## Supplementary figures and tables for "The individuality of shape asymmetries of the human cerebral cortex"

### Appendix 1

#### Figure Supplements

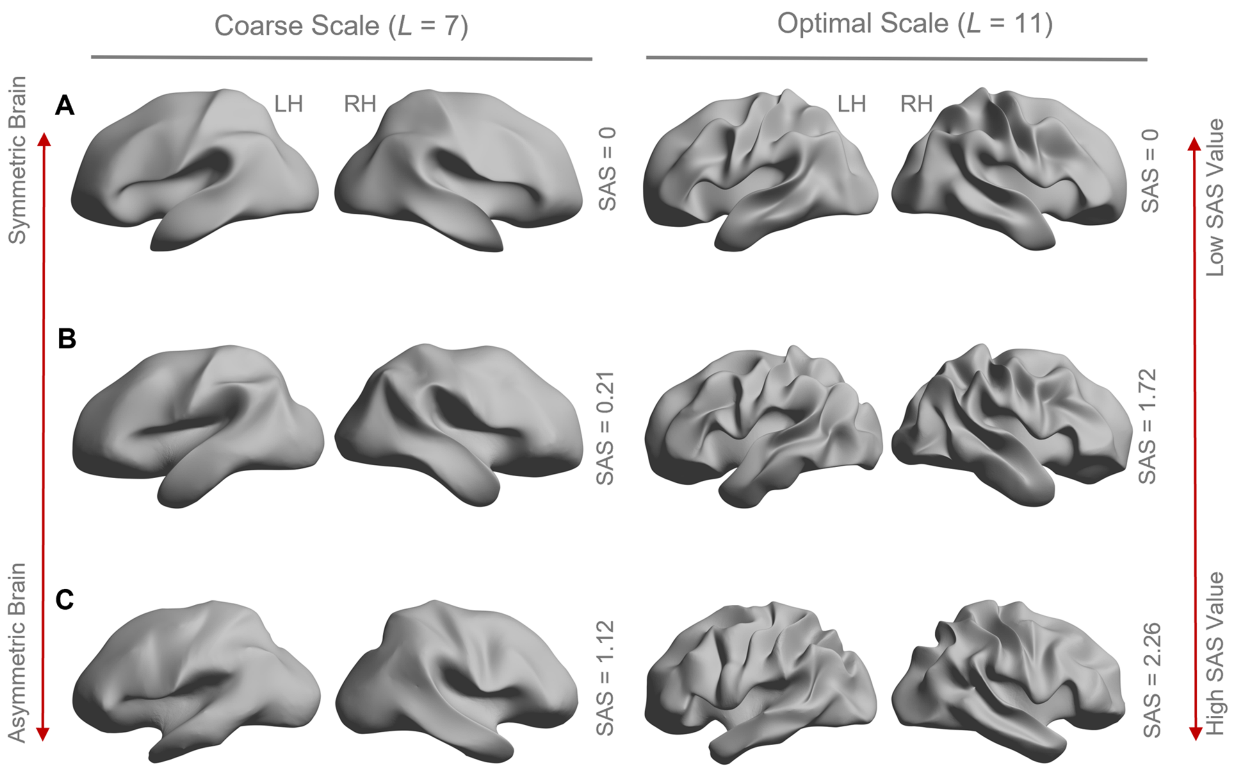


**Figure 1—figure supplement 1**. Higher SAS values characterize brains with stronger cortical shape asymmetries. Panels **(A)**, **(B)**, and **(C)** show left and right cortical surface reconstructions for three individuals showing varying levels of the SAS, from perfectly symmetric (panel A) to highly asymmetric (panel C). The left panel shows reconstructions at a coarse spatial scale corresponding to the first seven eigen-groups with a wavelength of about 55 mm. The right panel shows a reconstruction at the optimal scale for SAS identifiability, corresponding to the first 11 eigen-groups and a wavelength of about 37 mm. The perfectly symmetric brain in panel A was created by projecting the left hemisphere to the right hemisphere using the population-based template (fsaverage). The SAS value is zero for this case. The surfaces in panels (B) and (C) correspond to individual participants with moderate (B) and strong (C) asymmetry. The gradations

of asymmetry can be appreciated visually. As expected, the participant in panel C has a higher SAS than the participant in panel B. The SAS values shown here are the absolute mean values.

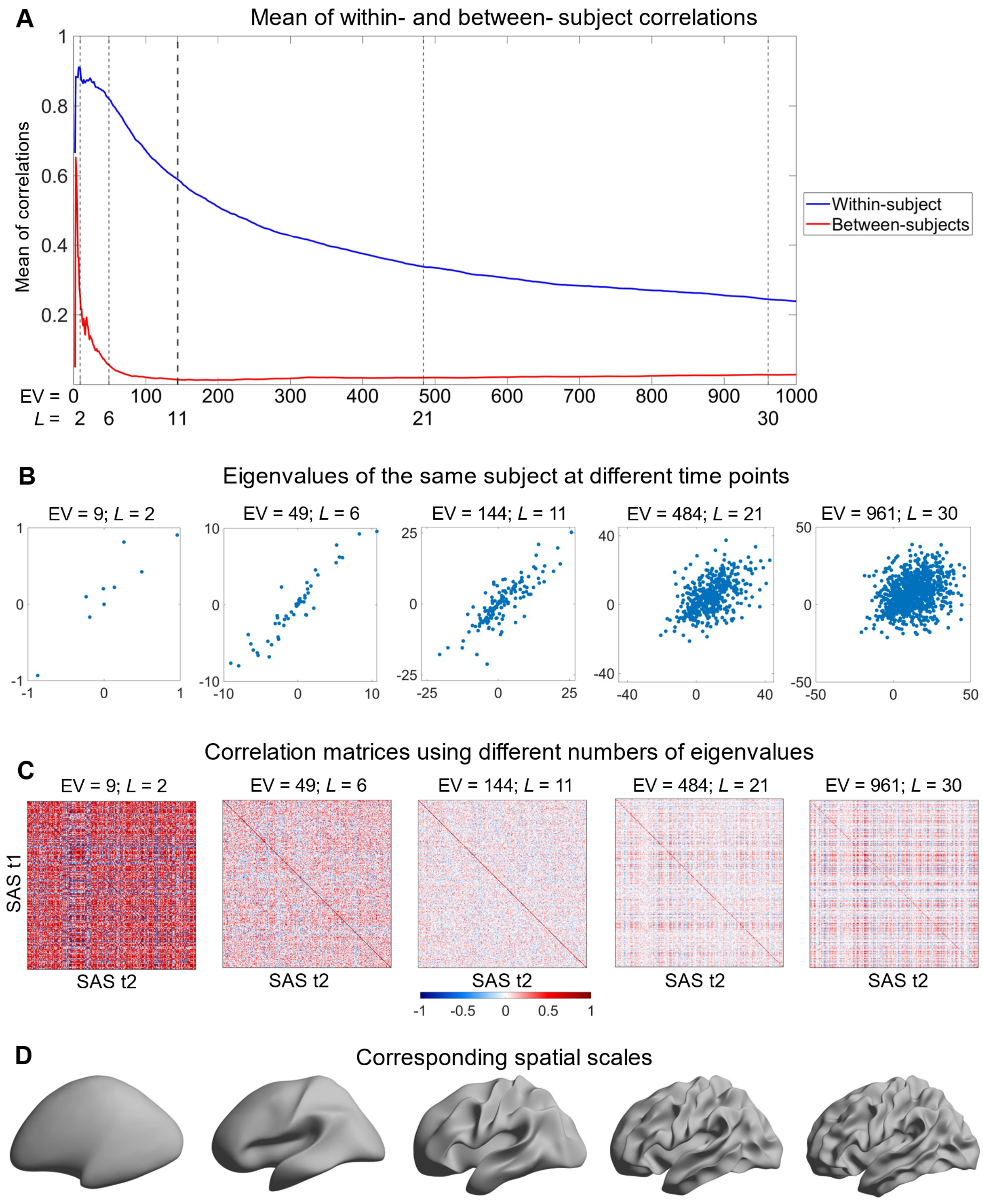


**Figure 2—figure supplement 1.** Understanding the identifiability score. Here, we use the shape asymmetry signatures from the OASIS-3 subjects (n = 233) as an example. **(A)** The mean of both within- and between- subject correlations decrease at finer scales, but the between-subject

correlations are lower and decline faster than the within-subject correlations. **(B)** The same subject at time 1 (t1) and time 2 (t2) with different numbers of eigenvalues. **(C)** Pearson correlation matrices using different numbers of eigenvalues. The diagonal elements are the within-subject correlations, and the off-diagonal elements are the between-subject correlations. Both within- and between- subject correlations are high from the very coarse scales (e.g., 9 eigenvalues in panel (C)); both correlations are low if fine scales are involved (e.g., 961 eigenvalues in panel (C)). The number of eigenvalues with peak identifiability score (144 eigenvalues; $L$ = 11) maximizes the difference between the between-subject and within-subject correlations. **(D)** Cortical surfaces reconstructed at spatial scales correspond to the eigen-groups in panel (C).

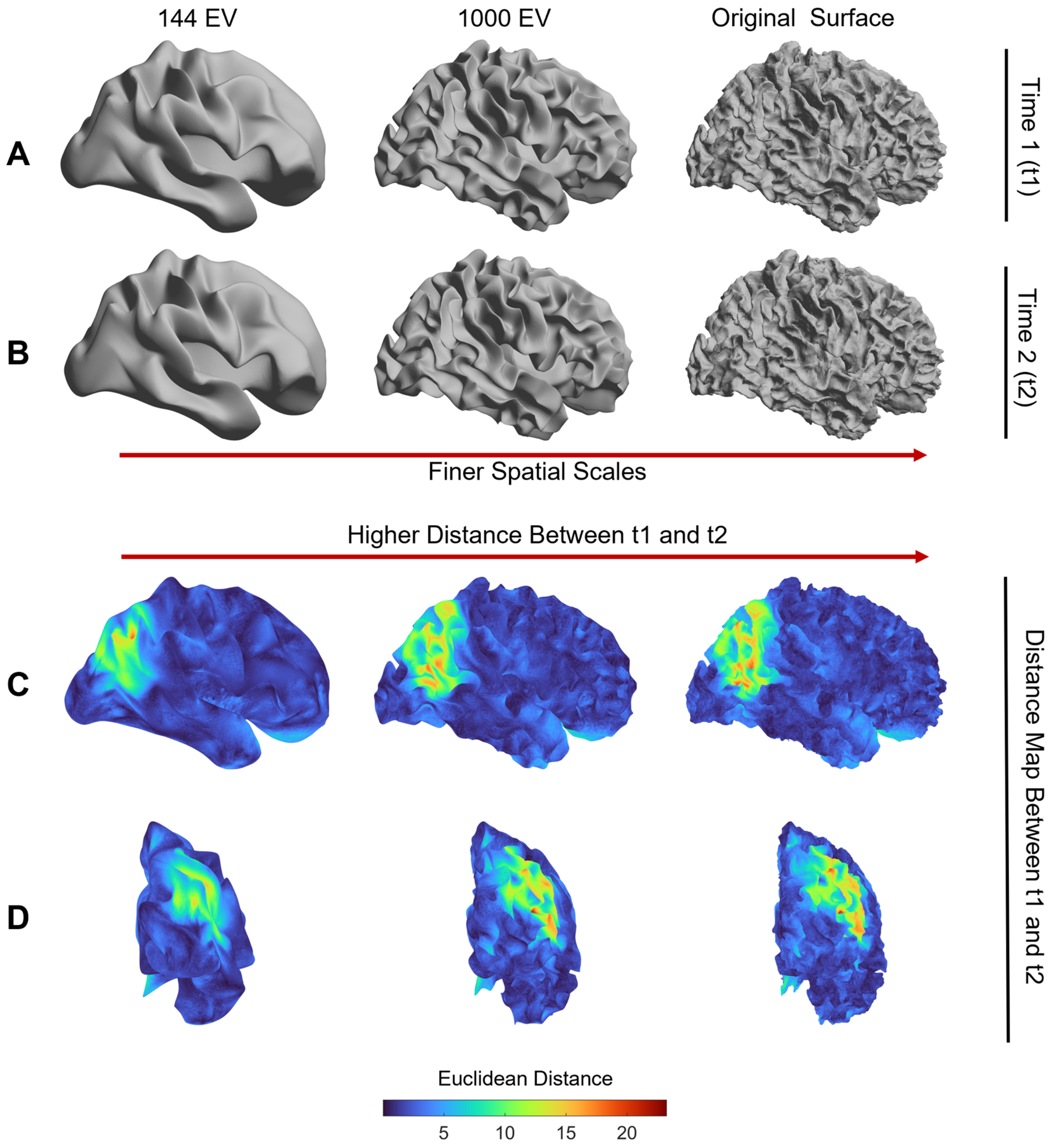


**Figure 2—figure supplement 2**. Inter-session variability in cortical shape is higher at more fine-grained spatial scales. Panels **(A)** and **(B)** show the white surface of one participant from the OASIS-3 dataset reconstructed at three spatial scales (i.e., using 144 eigenmodes, 1000 eigenmodes, and the full cortical surface) for time 1 and time 2 sessions, respectively. Panels **(C)** and **(D)** map the Euclidean distance of mesh vertices between time 1 and time 2 at each spatial

scale. The inter-session distances increase at finer scales (i.e., the original surface at the right panel). The images are registered on the fsaverage template.

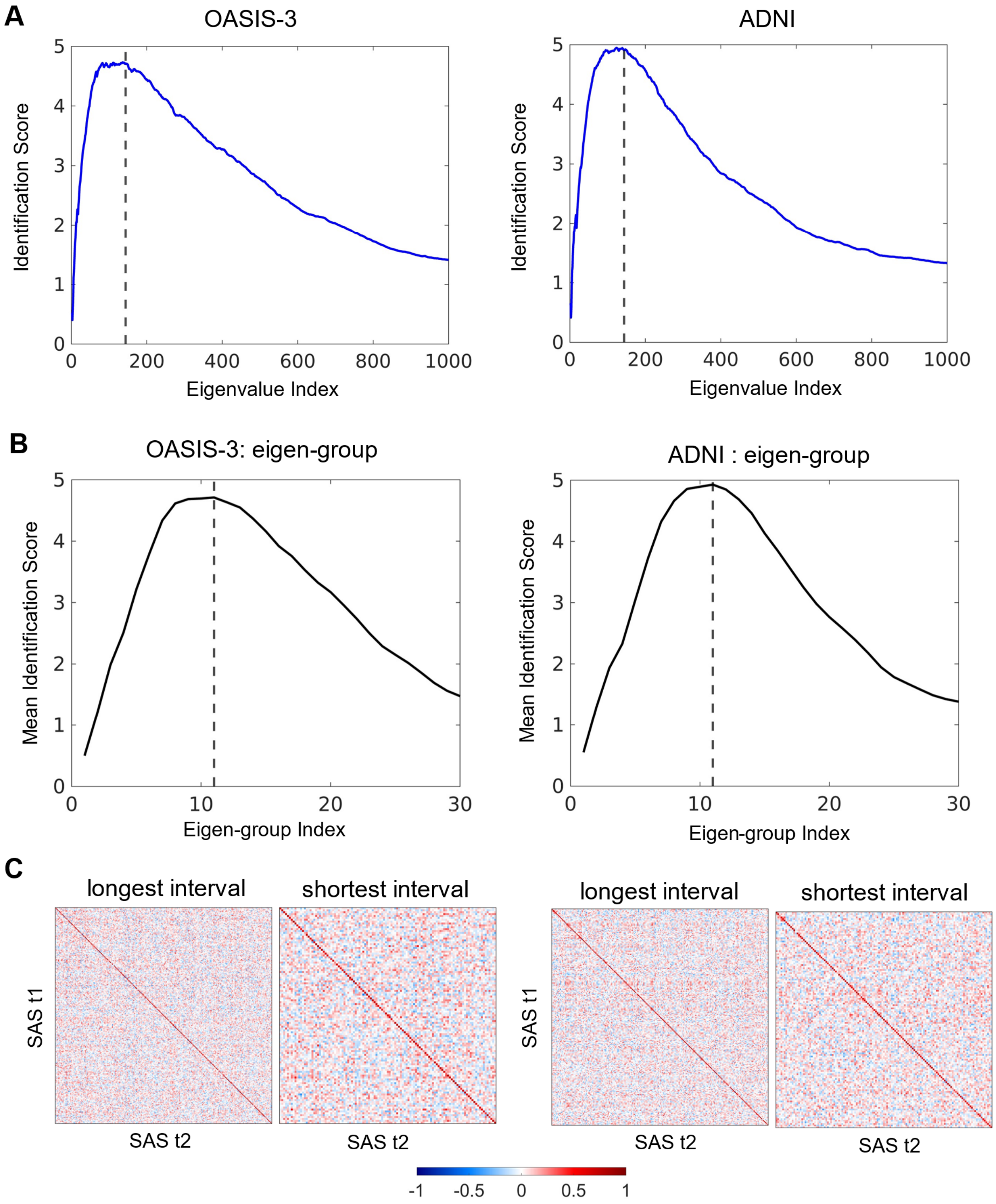


**Figure 2—figure supplement 3.** Subject identifiability scores re-calculated for data from MRI sessions with the longest inter-sessional interval. The optimal spatial scales determined by eigen-groups are identical to the initial analysis using the shortest inter-sessional interval. **(A)** The peak subject identifiability score occurs at the combination of the first 136 and 139 eigenvalues in the

OASIS-3 (n = 115) and ADNI (n = 135) data, respectively. **(B)** The peak mean subject identifiability score occurs at the first $11^{th}$ eigen-groups, representing the first 144 eigenvalues, in both the OASIS-3 and ADNI data. **(C)** Pearson correlation matrices calculated using the first 144 eigenvalues for the OASIS-3 and ADNI data by using both the shortest (OASIS-3: n = 233; ADNI: n = 208) and longest inter-sessional intervals.

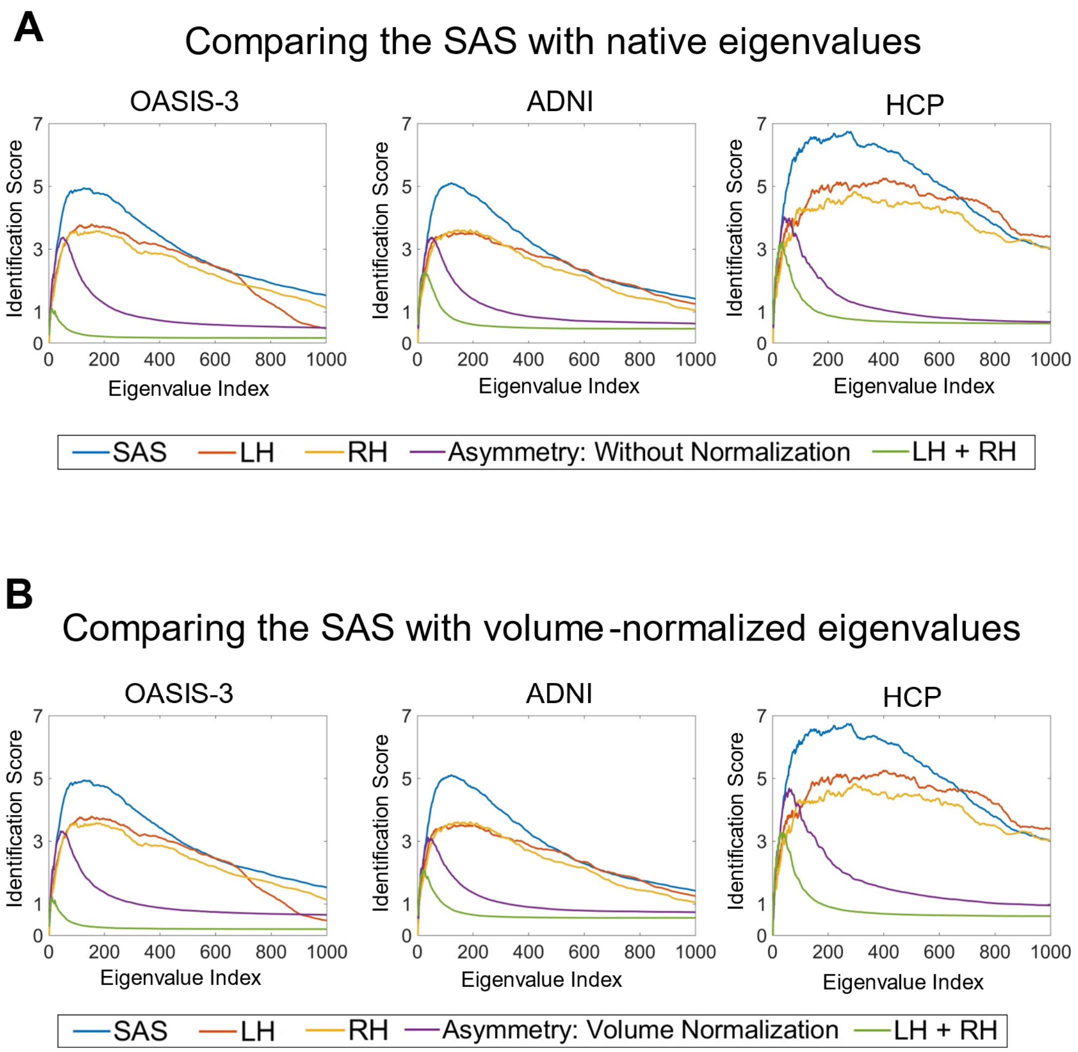


**Figure 3—figure supplement 1.** Comparing identifiability scores between the shape asymmetry signature (SAS) with either native eigenvalues or volume-normalized eigenvalues. The identifiability scores calculated from the surface area normalized SAS are generally higher than the scores calculated using native eigenvalues and eigenvalues with volume normalization (but without surface area normalization) for individual hemispheres, the combination of both hemispheres, and asymmetry across three datasets (OASIS-3: n = 233; ADNI: n = 208; HCP: n = 45). **(A)** identifiability scores calculated from native eigenvalues (except the blue lines, which are the SAS); **(B)** identifiability scores calculated from eigenvalues with volume normalization (except the blue lines, which are the SAS).

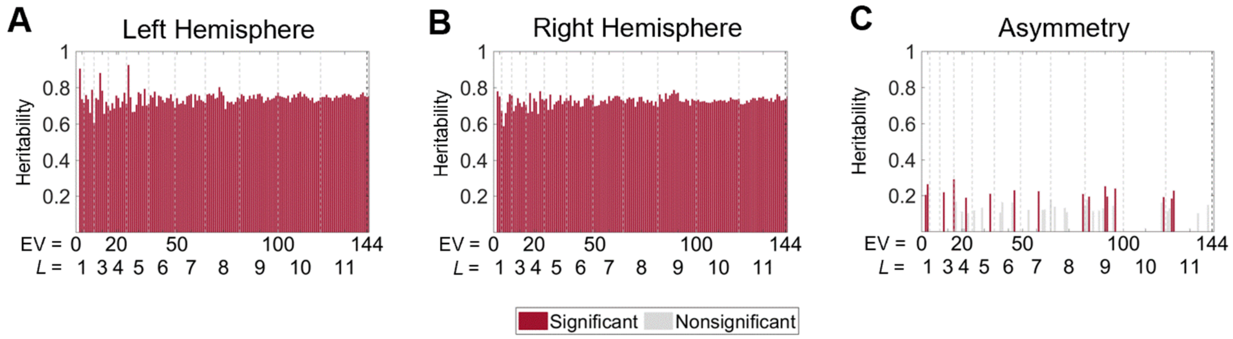


**Figure 6—figure supplement 1.** Heritability of cortical shape with volume normalization but without normalizing the surface area. **(A)** and **(B)** The heritability of the eigenvalues from the left (A) and right (B) hemispheres are uniformly high across all eigenvalues, and the scale-specific effects are eliminated. The heritability estimates are very close to the heritability of the mean of the cortical volumes across all regions of the MMP1 atlas ($h^2 = 0.77$ for the left hemisphere and $h^2 = 0.76$ for the right hemisphere). This result indicates that even normalizing the cortical volume, the heritability estimates are still highly influenced by the volume rather than purely by the shape. **(C)** Heritability estimates of the asymmetry are lower than that of the individual hemispheres but still have no scale effects. Statistical significance is evaluated after FDR-correction. We use 79 same-sex dizygotic twin pairs, 138 monozygotic twin pairs, and 160 of their non-twin siblings.

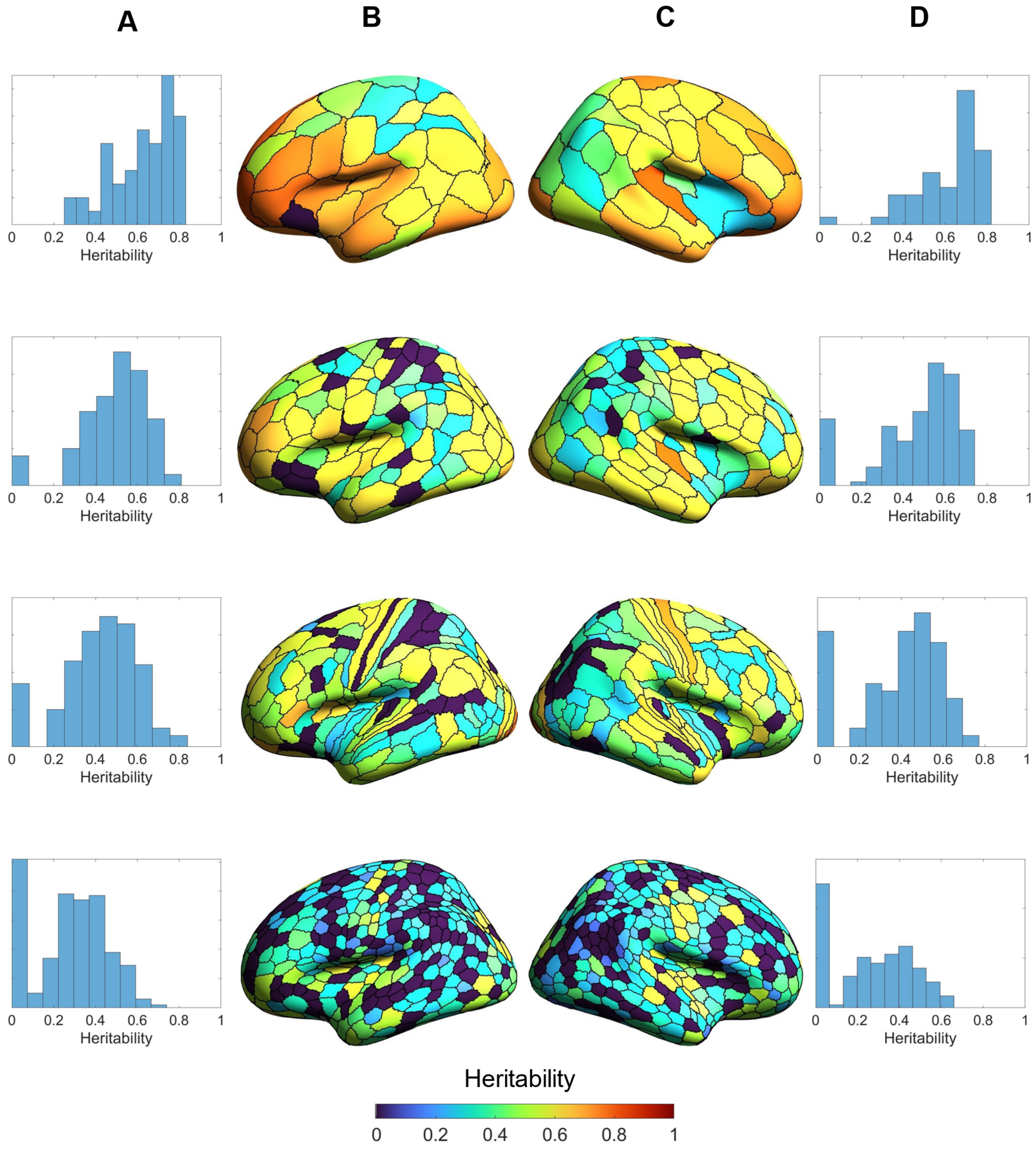


**Figure 6—figure supplement 2.** Heritability estimates of regional volumes of individual hemispheres across four parcellation resolutions: Schaefer 100, Schaefer 300, HCP-MMP1, and Schaefer 900 (top to bottom panels). Generally, heritability estimates are higher at coarser (upper panels) than finer parcellation resolutions (lower panels). **(A)** and **(D)** are the distributions of the

regional heritability estimates of the left (A) and right (D) hemispheres. **(B)** and **(C)** are heritability estimates of each region of the left (B) and right (C) hemispheres.

**Supplementary File 1**. Wavelength and eigenvalue indices of each eigen-group.

| Eigen-group Index | 0 | 1 | 2 | 3 | 4 |
|---|---|---|---|---|---|
| Eigenvalues | 1$^{st}$ | 2$^{nd}$–4$^{th}$ | 5$^{th}$–9$^{th}$ | 10$^{th}$–16$^{th}$ | 17$^{th}$–25$^{th}$ |
| Wavelength (mm) | N/A | 297.673 | 171.862 | 121.525 | 94.133 |
| Eigen-group Index | 5 | 6 | 7 | 8 | 9 |
| Eigenvalues | 26$^{th}$–36$^{th}$ | 37$^{th}$–49$^{th}$ | 50$^{th}$–64$^{th}$ | 65$^{th}$–81$^{st}$ | 82$^{nd}$–100$^{th}$ |
| Wavelength (mm) | 76.859 | 64.958 | 56.255 | 49.612 | 44.374 |
| Eigen-group Index | 10 | 11 | 12 | 13 | 14 |
| Eigenvalues | 101$^{st}$–121$^{st}$ | 122$^{nd}$–144$^{th}$ | 145$^{th}$–169$^{th}$ | 170$^{th}$–196$^{th}$ | 197$^{th}$–225$^{th}$ |
| Wavelength (mm) | 40.138 | 36.641 | 33.705 | 31.205 | 29.050 |
